# Effects of long-term warming and enhanced nitrogen and sulfur deposition on microbial communities in a boreal peatland

**DOI:** 10.1101/704411

**Authors:** Magalí Martí, Alexander Eiler, Moritz Buck, Stefan Bertilsson, Waleed Abu Al-Soud, Søren Sørensen, Mats B. Nilsson, Bo H. Svensson

## Abstract

With ongoing environmental change, it is important to understand ecosystem responses to multiple perturbations over long time scales at *in situ* conditions. Here, we investigated the individual and combined effects of 18 years of warming and enhanced nitrogen and sulfate deposition on peat microbial communities in a nutrient-poor boreal mire. The three perturbations individually affected prokaryotic community composition, where nitrogen addition had the most pronounced effect, and its combination with the other perturbations led to additive effects. The functional potential of the community, characterized by shotgun metagenomics, was strongly affected by the interactive effects in the combined treatments. The responses in composition were also partly reflected in the functional gene repertoire and in altered carbon turnover, i.e. an increase of methane production rates as a result of nitrogen addition and a decrease with warming. Long-term nitrogen addition and warming-induced changes caused a shift from *Sphagnum*-dominated plant communities to vascular plant dominance, which likely transact with many of the observed microbial responses. We conclude that simultaneous perturbations do not always lead to synergistic effects, but can also counteract and even neutralize one another, and thus must be studied in combination when attempting to predict future characteristics and services of peatland ecosystems.

## Introduction

Despite key roles of microbial communities in controlling carbon and nutrient dynamics within ecosystems, few studies have addressed effects of multiple drivers (e.g. anthropogenic stressors) on microbial taxonomic diversity and the associated genome-encoded traits underpinning realized ecosystem services (cf. Castro et al. 2010, Li et al. 2013, Andresen et al. 2014, Shen et al. 2014, Contosta et al. 2015). Given the multitude of human-derived disturbances, including climate change and atmospheric deposition of anthropogenic emissions, it is important to understand microbial community responses to multiple and parallel changes in the environment. Microorganisms will potentially react to such external impacts by shifting their community composition and function. This may lead to ecosystem structure repercussions causing feedbacks on e.g. the climate system via changes in greenhouse gas turnover and other ecosystem-scale processes. However, altered ecosystem services resulting from microbial community shifts may also be attenuated by functional redundancy within the community, assuming that many species are able to mediate the same functions in the ecosystem (Lawton & Brown 1993, Nielsen et al. 2011, Tully et al. 2018).

Microbial metabolism is central for most ecosystem services due to its central role in the turnover of essential nutrients and biogeochemical cycles (Martiny et al. 2015). For example, changes in the abundance of key microbial taxa able to produce or oxidize methane will influence the rate of methane emissions from various ecosystems that feature relevant redox conditions (McCalley et al. 2014). It is now tractable to characterize such genome-encoded functional traits with community-scale metagenome sequencing.

*In situ* manipulation experiments are well suited to test the responses of ecosystems and their microbial communities to various disturbance scenarios. Except for long-running trials in agricultural and forestry systems (Ramirez et al. 2010, Fierer et al. 2012, Leff et al. 2015, Zhou et al. 2015, Boot et al. 2016, Zeng et al. 2016), generally, *in situ* manipulations of soil ecosystems have mainly been conducted over short-to intermediate-length time periods, e.g. between 1 to 5 years. Thus, several such short time field studies on effects of nitrogen amendment and warming on diversity, function and abundance of microbial communities have been reported (Castro et al. 2010, Li et al. 2013, Andresen et al. 2014, Shen et al. 2014, Contosta et al. 2015). Yet, to identify environmental responses over ecologically relevant timescales, while at the same time accounting for short-term disturbance effects, long-term (at least decadal) field experiments are needed (Rinnan et al. 2007, Eriksson et al. 2010a, Contosta et al. 2015). Moreover, as all ecosystems are influenced by multiple stressors, we need to understand how interactive effects of multiple effectors (synergistic or antagonistic perturbations) influence microbial communities if we are to robustly predict ecosystem responses to global warming, changes in atmospheric nutrient deposition and anthropogenic pollution in general.

Ever since the last de-glaciation, northern peatlands have played an important role in the global carbon balance, and are currently estimated to hold 30% of the global soil carbon, i.e. carbon stocks estimated at 270 - 600 Pg of organic C (Gorham 1991, Turunen et al. 2002, Yu 2012). Due to slow rates of microbial decomposition, there is an imbalance between primary production and degradation in these ecosystems. Thus, undisturbed peatlands are generally contemporary net sinks of carbon dioxide (CO2), while at the same time being significant sources of methane (CH4) to the atmosphere (Roulet et al. 2007, Nilsson et al. 2008, Turetsky et al. 2014). Therefore, the impact of the predicted global climate change and atmospheric nitrogen (N) and sulfate (S) deposition on the peatland net ecosystem carbon balance (NECB) is a topic of major concern (Granberg et al. 2001, Galloway et al. 2004, Gauci et al. 2004, Phoenix et al. 2012).

Increased N availability is known to strongly affect plant species composition and performance of different plant functional types in peatlands (Damman 1988, Bridgham et al. 1996, Eppinga et al. 2010, Limpens et al. 2011) as N is often a limiting nutrient (Wang & Moore 2014). This is clearly demonstrated by the shift towards vascular plant-dominated vegetation in the long-term high N deposition experiment at Degerö Stormyr (Wiedermann et al. 2007, Eriksson et al. 2010a, Eriksson et al. 2010b). As a result, methane emissions increased in response to the enhanced N deposition (Eriksson et al. 2010b), whereas warming led to a decrease in methane production, oxidation and emissions (Eriksson et al. 2010a, Eriksson et al. 2010b).

It has been argued that novel mechanistic insights on biogeochemical dynamics can be obtained by studying peat microbial community composition and function, and that such information can further improve predictions of ecosystem responses to global change (Bragazza et al. 2015). To investigate the isolated and interactive effects of warming, N and S deposition on the peat microbial community and function, we applied high throughput sequencing approaches to peat samples collected after 18 years of continuous *in situ* field manipulation. We took advantage of a full factorial experimental design that was established at Degerö Stormyr in 1995, consisting of nitrogen (NH_4_NO_3_) and sulfate (Na_2_SO_4_) amendments and warming (plots scale green-house covers) simulating the predicted effect of climate change (Granberg et al. 2001). Overall, simulated increased N deposition had the most pronounced effect on bacterial as well as archaeal communities. Multiple stressors interacted to give responses at the level of taxonomic and functional diversity, which influenced the functional potential of the ecosystem with regards to methane production, sulfate reduction, nitrate reduction and polymer hydrolysis. Thus, our experiment emphasizes the need to study the effects of climate change on microbial communities in the context of multiple environmental changes and anthropogenic-induced perturbations.

## Materials and Methods

### Site description and peat samples collection

Peat samples were collected from a full factorial experimental design in a *Sphagnum-dominated* oligotrophic area at Degerö Stormyr, in North Sweden (64°11’N, 19°33’E, altitude 270 m above sea level). Briefly, the experimental site is a boreal oligotrophic minerogenic mire, with a surface water pH of ~4.5. The climate of the reference period (1961-1990) was characterized by a mean annual precipitation of 523 mm, a mean annual temperature of 1.2°C, and a mean July and January temperatures of 14.7°C and −12.4°C, respectively. Average weather conditions during 2001-2012 were as follows (Peichl et al. 2014): annual and growing season air temperatures 2.3°C and 11°C, respectively, annual and growing season precipitation of 666 and 395 mm, respectively, and the growing season mean water table level at 14 cm below peat surface. The dominant vascular plants are *Eriophorum vaginatum* (L), *Andromeda polifolia* (L). and *Vaccinium oxycoccos (L.).* The dominant moss species are *Sphagnum balticum* (Russ) C. Jens. and, *S. lindberghii* (Schimp). The experiment was established in 1995 (Granberg et al. 2001) and was conducted according to a full factorial experimental design including two levels of nitrogen (N) i.e. ambient (low-level) at 2 kg N ha^−1^ yr^−1^ and amendment (high-level) of NH_4_NO_3_ to reach a deposition at a level of 30 kg N ha^−1^ yr^−1^, two levels of sulfur (S) i.e. ambient at 3 kg S ha^−1^ yr^−1^, and amendment of Na_2_SO_4_ to a level of 20 kg S ha^−1^ yr^−1^, and two levels of greenhouse (GH) treatment, i.e. high level GH with a transparent cover or ambient (low level) of GH i.e. without a cover. Each experimental combination was performed in duplicate. The elevated levels of N and S correspond to the annual deposition amounts in southwest Sweden at the time for the start of the experiment. For a detailed description of the site and experimental design see Granberg et al. (2001), and for details on treatment effects on vegetation composition see Wiedermann et al. (2007). One peat core (0-40cm) was collected from each field plot (n=16) on August 14^th^ and 15^th^ 2013. From each core, 5 cm^3^ of peat were subsampled at five depths from below the *Sphagnum* surface: 7-11 cm (A), 11-15 cm (B), 15-19 cm (C), 19-23 cm (D) and 23-27 cm (E). The samples were stored in 50 mL sterile tubes containing 2 mL of LifeGuard™ Soil Preservation Solution (MoBio Laboratories, Hameenlinna, Finland), and kept at room temperature (<20°C) for at most 24 hours before freezing at −20°C.

### Sample preparation

Before extraction, peat samples were thawed over night at 4°C and centrifuged at 2500 x g for 5 min. Total RNA and DNA were co-isolated from 2 g wet peat and recovered in 50 μl using the RNA PowerSoil^®^ Total RNA Isolation Kit together with the RNA PowerSoil^®^ DNA Elution Accessory Kit (MoBio Laboratories), according to manufacturer’s instructions. Extract concentrations were measured with the Quant-iT RNA HS assay and the Quant-iT dsDNA HS assay kits together with the Qubit fluorometer (Invitrogen, Lidingö, Sweden). The RNA and DNA extraction yields ranged 2.5 – 122 ng μl^−1^ and 0.5 – 600 ng μl^−1^, respectively. Two extractions from each sample were performed and combined after extraction. RNA combined extractions were concentrated using the RNA Clean & Concentrator™-25 (Zymo Research, Taby, Sweden), in accordance with the manufacturer’s instructions. DNA was concentrated by the use of a Vacufuge^®^ vacuum concentrator 5301 (Eppendorf, Horsholm, Denmark). DNA residues were removed from the concentrated RNA extracts by digestion using 2U TURBO DNase (Ambion-Life Technologies, Stockholm, Sweden) for 1 h at 37°C and according to manufacturer’s instructions. Reverse transcription was performed adding 2 μl of DNAse-treated RNA to 17 μl reaction mixture containing 1X Expand Reverse Transcriptase Buffer (Roche Diagnostics, Mannheim, Germany), 10mM of Dithiothreitol (DTT) solution (Roche diagnostics), 5 mM of dNTPs (New England BioLabs Inc., Glostrup, Denmark) and 250 nM of random hexamers (TAG Copenhagen A/S, Copenhagen, Denmark). After 2 min incubation at 42°C in a DNA engine DYAD™ Peltier Thermal Cycler (MJ researchBio-Rad Laboratories, Hercules, CA), 1 μl of Expand Reverse Transcriptase (Roche Diagnostics) was added to the mixture and incubated at 42°C for 40 min followed by 30 min at 50°C and 15 min at 72°C. RNA template addition was in the range of 8 – 300 ng.

### Amplicon sequencing and sequences analysis

PCR amplification of the bacterial and archaeal 16S rRNA and 16S rRNA gene V4 region fragment was performed using the primer pair 515F/806R (Caporaso et al. 2012) for all samples (Table S1). The primers used for amplicon sequencing of the 16S rRNA gene were selected following the Earth Microbiome Project recommendation (http://www.earthmicrobiome.org/emp-standard-protocols/) representing the best choice at the initiation of the study. The primer 806R was previously modified to cover most available sequences in Genbank by Sundberg et al. (2013). More recently, these primers have been shown to be biased against Crenarcheota (Hugerth et al. 2014). Three μl of template (cDNA or DNA) were added to a 20 μl-reaction mixture, consisting of 1X AccuPrime PCR Buffer II (Invitrogen), 0.2 U of AccuPrime *Taq* DNA Polymerase High Fidelity (Invitrogen) and 500 nM of each primer. The PCR assay was performed in a DNA engine DYAD™ Peltier Thermal Cycler (Bio-Rad Laboratories) with the following conditions: 94°C for 2 min, followed by 35 cycles of 94°C for 20 s, 56°C for 30 s and 68°C for 40 s, and a final extension at 68°C for 5 min. PCR products were visualized on a 1% agarose gel. A second PCR using the same primers including adapters and indexes was performed under the same conditions as the previous PCR with the number of cycles reduced to 15. PCR amplicons were purified using Agencourt AMPure XP (Agencourt Bioscience Corporation, MA, USA) and concentration was measured using PicoGreen assay according to manufacturer’s protocol (Invitrogen). To ensure equal representation of each sample, 50 ng of each 16S rRNA and rRNA gene purified sample amplicons were pooled together before sequencing. Then the pooled mixture was purified and concentrated using the DNA Clean & Concentrator™-5 (Zymo Research) followed by quantification with the Quant-iT dsDNA HS assay kit and the Qubit fluorometer (Invitrogen). A paired-end 250-bp sequencing run was performed using the Illumina MiSeq instrument according to the MiSeq™ Reagent Kit v2 Preparation Guide (Illumina, Inc., San Diego, CA, USA).

Raw read data was processed using the ILLUMITAG pipeline (Sinclair et al. 2015). In short, the read pairs from the 16S rRNA gene and 16S rRNA were demultiplexed and joined using the PANDAseq software v2.4 (Masella et al. 2012). Next, assembled reads (from here referred to as “reads”) that did not have the correct primer sequences at the start and end of their sequences were discarded. Reads were then filtered based on their PHRED scores. Chimera removal and OTU (operational taxonomic unit) clustering at 1% sequence dissimilarity was performed by pooling all reads from all samples (but separately for genomic 16S rRNA and 16S rRNA) together and applying the UPARSE algorithm v7.0.1001 (Edgar 2013). Here, any OTU containing less than two reads was discarded. Each OTU was subsequently taxonomically classified by operating a similarity search against the SILVAmod databases and employing the CREST assignment resource (Lanzen et al. 2012). Finally, plastid and mitochondrial OTUs were removed.

We obtained a total of 4 965 761 sequence reads (including both, the 16S rRNA and 16S rRNA gene), from which 4 811 127 sequences belonged to the domain Bacteria and 15 444 to the domain Archaea. Bacterial and archaeal sequences were analysed together as the combined prokaryote community. Prior to estimating alpha diversity, to minimize the impact of varying sequencing depth among the samples, the reads were rarefied to 5315 and 5455 sequences per sample for RNA and DNA, respectively. Three RNA and 2 DNA samples were excluded due to low number of sequences (Table S1). Prokaryote richness and evenness (alpha diversity) were estimated using the Chao1 and Pileou’s indices, where a Pielou index of 1 represents absolute evenness. Prior to beta-diversity analyses and to avoid excluding samples, the number of reads for individual samples were rarefied to the minimum number of sequences observed per sample: 3331 and 3348 sequences for 16S rRNA and the 16S rRNA gene, respectively.

### Shotgun metagenome sequencing, assembly and annotation

Shotgun sequencing was performed on samples from depths B (11-15 cm) and D (19-23 cm), (Table S1). Depth B corresponds to the level around which the mean growing season water table occurs, which is the depth horizon with the most metabolic activity. Depth D is below the growing season water table level fluctuations and, thus, continuously anoxic. 1ng of each DNA sample (from B and D depths) were used for tagmentation using the Nextera XT (Illumina, Inc., San Diego, CA, USA), according to manufacturer’s instructions. Tagmented samples were purified using Agencourt AMPure XP (Agencourt Bioscience Corporation, MA, USA). Purified samples were visualized on a 1% agarose gel to ensure the libraries range was within 300-1000 bp. Libraries concentrations were measured with Quant-iT dsDNA HS assay and the Qubit fluorometer (Invitrogen). To ensure equal representation of each sample, 10 ng of each sample library were pooled together before sequencing. A paired-end 150-bp sequencing run was performed using the Illumina Rapid Run on an Illumina HiSeq 2500 platform (Illumina, Denmark), according to manufacturer’s instructions.

The obtained reads were quality-trimmed using Sickle (Joshi & Fass 2011), and all samples were co-assembled with a range of k-mer values (from 31 to 101 with increments of 10) using Ray (Boisvert et al. 2012). The resulting assemblies were subsequently fragmented *in silico* into successive sequences of 2000 base pairs overlapping by 1900 bp and were then merged using 454 Life Sciences’s software Newbler (Roche, Basel, Switzerland) as previously described (Hugerth et al. 2015). The clean reads of all samples were mapped to the merged assembly using Bowtie (Langmead & Salzberg 2012) after processing with SAMtools (Li et al. 2009). Duplicates were removed using Picard (Broad Institute, Cambridge, MA, USA). Finally, coverage was computed using BEDTools (Quinlan & Hall 2010). PFAM (protein families) annotation was performed using HMMer (Eddy 2011) using the PFAM A database (Finn et al. 2014). For each sample, the coverage of all detected PFAMs was normalized by dividing it by the mean coverage of a set single copy PFAMs (Rinke et al. 2013) in order to compile the coverage in a “per genome equivalent” form. PFAM tables standardized to genome equivalents were resampled by removing PFAMs with smaller genome equivalents than the highest minimum genome equivalent (6.9×10^−4^) of sample NxGH.

### Statistical analyses

The multifactorial experiment consists of three treatment factors at two levels (2^3^-design) with field duplicates for each treatment. Thus, the statistical evaluation is based on n=8 for the main factors N, S and GH and n=4 for the 2-way interaction treatments (NxS, NxGH and SxGH) and n=2 for the 3-way interaction (NxSxGH), see Table S2 for the treatment effect evaluation matrix. After 10 years of treatment, the addition of N had significantly reduced the distance between the mire surface and the growing season average water table (Eriksson et al. 2010b). To account for this gradual change and the inherent variation among the plots, the sampled depths were classified according to their positions relative to the average growing seasonal water table level within each plot as given by (Eriksson et al. 2010a), (Table S3). This resulted in three different depth horizons: an upper layer above the growing season mean water table level, which will be the most oxic of the three (AWT), a layer around the growing season mean water table level (WT) and a third layer below the growing season mean water table level characterized by permanent anoxic conditions (BWT).

ANOVA was applied to test for the effects of the treatments on the prokaryotic alpha-diversity. Permutation analysis of variance (PERMANOVA) was applied to test the hypothesis that the prokaryotic community composition and its functional potential differed among the treatments. To assess whether the microbial composition turnover in the treatment plots followed the same direction, ordination by non-metric multidimensional scaling was used. In order to understand how the response of the microbial community was distributed across the different phyla, we used the integrated occurrence of each phylum along the entire peat profiles and calculated the average change in their relative abundance derived from the 16S rRNA for the high treatment levels in relation to their corresponding low levels. The phyla were sorted according to their relative abundance in the combined 16S rRNA dataset and are presented with either positive or negative responses to the treatments. In this context, it should be noted that changes at the phylum level might only be a rough representation of changes in the functional capacity of the communities. To assess individual metabolic traits in the different treatments, we applied generalized linear models (GLMs) on genome equivalent standardized PFAMs across the different treatments. We focused on PFAMs predicting key enzymes involved in the anaerobic degradation of soil organic matter, as well as methanotrophy (Table S4). The resulting differentially abundant categories (taxa or functional subsystems) among samples were identified based on p<0.05 and false discovery rate was estimated (Benjamini & Hochberg 1995). A distance-based redundancy analysis (db-RDA) was applied to explore possible multiple linear correlations between the microbial community composition and the vegetation composition previously reported (Eriksson et al. 2010b). Correspondence between the different data sets was investigated using procrustes superimposition combined with a randomisation test (Peres-Neto & Jackson 2001). Bray-Curtis distance was used when a distance matrix was required applying 999 permutations. The statistic discrimination throughout the analyses was at a significance level of 0.05.

### Nucleotide sequence accession numbers

The sequence data generated in this study was deposited to the NCBI Sequence Read Archive and is accessible through accession number PRJEB14741.

## Results

### Microbial community composition

High throughput sequencing data was used to follow prokaryotic community composition and metabolic traits responding to the perturbations in the factorially designed experiment. High concordance was observed between the 16S rRNA gene and 16S rRNA-derived community compositions, as determined by procrustes superimposition (p = 0.001, R = 0.9; Table S5). Comparisons of treatments revealed that the three main factors (nitrogen (N), sulfate (S) and warming (GH)) significantly affected the prokaryote community compositions (i.e. the beta-diversity; Table 1). In all cases, enhanced inorganic N deposition (high level) significantly affected the community composition at all depth horizons, contributed the most to the explained variance (Table 1), and showed the highest degree in dissimilarity compared to the ambient N (low level) (Fig. 1). In contrast to beta-diversity describing compositional changes (as determined by Bray-Curtis distances), alpha-diversity assessed as prokaryotic richness and evenness estimates, revealed only few significant responses to the perturbations (Table 1). Richness increased with the warming effect, while evenness overall decreased as a result of all the perturbations (Fig. S1).

**Fig. 1.**
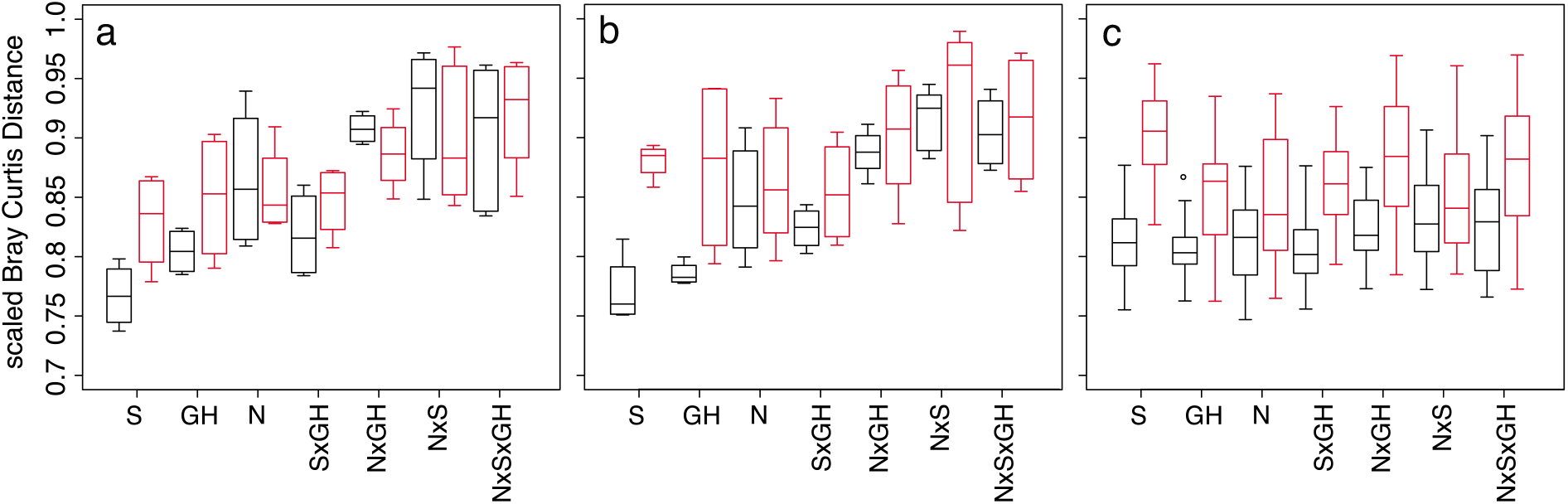
Prokaryote community composition (β-diversity) turnover in relation to the low treatment levels. a: above the growing season mean water table level. b: around the growing season mean water table level. c: below the growing season mean water table level. Black: community composition derived from the 16S rRNA gene. Red: community composition derived from the 16S rRNA. A Bray-Curtis distance of 0 indicates a complete overlap in community composition between the high and low treatment levels, while a Bray-Curtis distance of 1 indicates complete dissimilarity. Note that the scale on the y-axes starts at 0.7. The number of replicates for the main, two- and three-way interaction effects are 8, 4 and 2, respectively.

**Table 1.** Statistical tests for the 16S rRNA gene sequences, 16S rRNA sequences and protein families derived from the metagenome (PFAM) in relation to the treatments and the standardized depths: above the growing season water table (AWT), around the growing season mean water table (WT), and below the growing season mean water table (BWT). PFAMs were only analysed at the AWT and BWT depths. N, S and GH refer to the main treatment effects of nitrogen, sulfur and greenhouse, and NxS, NxGH and SxGH refer to their two-way interactions, while NxSxGH represents the three-way combination. Only R2-values from the significant results of a permutational multivariate analysis of variance (PERMANOVA) are presented. Similar for the alpha-diversity, only Pielou (evenness index) and Chao (richness index) are represented from the significant results of an ANOVA.

| Data |  | Effect | Standardized depth |  |  |
| --- | --- | --- | --- | --- | --- |
|  |  |  | AWT | WT | BWT |
| Beta-diversity | 16S rRNA gene | N | 0.25*** | 0.18** | 0.08*** |
|  |  | S | ns | 0.06* | 0.03* |
|  |  | GH | 0.07* | 0.09** | 0.05*** |
|  |  | NxS | ns | 0.05* | ns |
|  |  | NxGH | 0.06* | 0.05* | ns |
|  |  | SxGH | ns | ns | 0.03* |
|  |  | NxSxGH | 0.05* | ns | 0.03* |
|  | 16S rRNA | N | 0.15** | 0.12* | 0.09*** |
|  |  | S | ns | 0.07* | 0.04** |
|  |  | GH | ns | 0.06* | 0.05** |
|  |  | NxS | ns | 0.05* | 0.03* |
|  |  | NxGH | ns | ns | ns |
|  |  | SxGH | ns | ns | ns |
|  |  | NxSxGH | ns | ns | ns |
| Functional potential | PFAM | N | ns |  | 0.14* |
|  |  | S | ns |  | ns |
|  |  | GH | ns |  | ns |
|  |  | NxS | ns |  | ns |
|  |  | NxGH | ns |  | ns |
|  |  | SxGH | ns |  | 0.13* |
|  |  | NxSxGH | ns |  | ns |
| Alpha-diversity | 16S rRNA gene | N | Pielou* | Chao*,Pielou** | ns |
|  |  | S | ns | Chao** | Chao**;Pielou* |
|  |  | GH | ns | ns | Chao* |
|  |  | NxS | ns | Pielou* | ns |
|  |  | NxGH | ns | ns | ns |
|  |  | SxGH | ns | Chao**;Pielou** | ns |
|  |  | NxSxGH | ns | ns | Pielou* |
|  | 16S rRNA | N | ns | ns | ns |
|  |  | S | ns | ns | Chao*;Pielou* |
|  |  | GH | ns | ns | ns |
|  |  | NxS | ns | ns | ns |
|  |  | NxGH | ns | ns | ns |
|  |  | SxGH | ns | ns | ns |
|  |  | NxSxGH | ns | ns | ns |

The combined perturbations (i.e. high levels of NxS, NxGH, SxGH and NxSxGH) caused significant interactive effects that shifted the prokaryotic community (i.e. the beta-diversity: Table 1 and Fig. 1). For example, the 16S rRNA gene-derived prokaryotic community (beta-diversity) response to warming increased with enhanced N deposition in the two upper horizons, while enhanced S deposition significantly increased the warming effect below the water table. Such synergistic effects were also observed when combining N and S deposition. While treatment interactions appeared to be additive as based on the observation that the dissimilarity among communities increased when applying multiple perturbations, their directionality were not consistent (Fig. 1 and Fig. S2). This implies that communities experiencing impacts from multiple perturbations do not merely change in a linear fashion based on combinations of individual perturbations, but are emerging in response to the establishment of unique communities for a particular combination of effectors. For example, enhanced N deposition combined with another perturbation (NxS, NxGH and NxSxGH) tended to cause greater community changes compared to the single perturbations at the two upper layers (Fig. 1a and 1b). Below the growing season mean water table (BWT) horizon, the amplitudes of the community composition responses in comparison to the control were similar for all perturbations (Fig. 1c).

There was concordance between beta diversity of the plant vegetation, derived from Wiedermann et al. (2007), and the peat prokaryote community at all three depth horizons, as revealed by procrustes superimposition (p = 0.001, R = 0.4 - 0.7; Table S5). In addition, db-RDA analyses revealed that the prokaryotic composition in the treatments receiving NH_4_NO_3_ (high N) were positively correlated with the relative abundance of the dominant vascular plants *(E. vaginatum, A. polifolia and V. oxycoccus)* and negatively correlated with total *Sphagnum* and *S. balticum* coverage (Fig. S3).

Bacterial and archaeal 16S rRNA gene sequences and expressed 16S rRNA sequences were classified into 31 and 33 phyla, respectively. The main phyla were Verrucomicrobia and Acidobacteria accounting respectively for 39% and 26% of the reads in the 16S rRNA gene-derived community, while Acidobacteria accounted for 36% and Proteobacteria for 23% of the reads in the 16S rRNA-derived community. The abundance of the different phyla was clearly and differentially affected by the perturbations (Fig. 2). For example, the phyla Acidobacteria, Firmicutes, and Armatimonadetes decreased in relative abundance in response to all the treatments, while Chlorobi, Euryarchaeota, Fibrocateretes, and Tenericutes showed the opposite response. Verrucomicrobia and Actionabacteria increased in response to the main N effect and decreased in the other treatments, while candidate phylum BD1-5 had the opposite response. Cyanobacteria and Fusobacteria, increased in response to the main warming effect while for all the other treatments decreased and disappeared, respectively. Dictyoglomi increased with N, GH and NxGH while disappeared with all the other treatments. Thaumarchaetoa and candidate division TM7 disappeared with warming while increased with N, S and NxS perturbations.

**Fig. 2.**
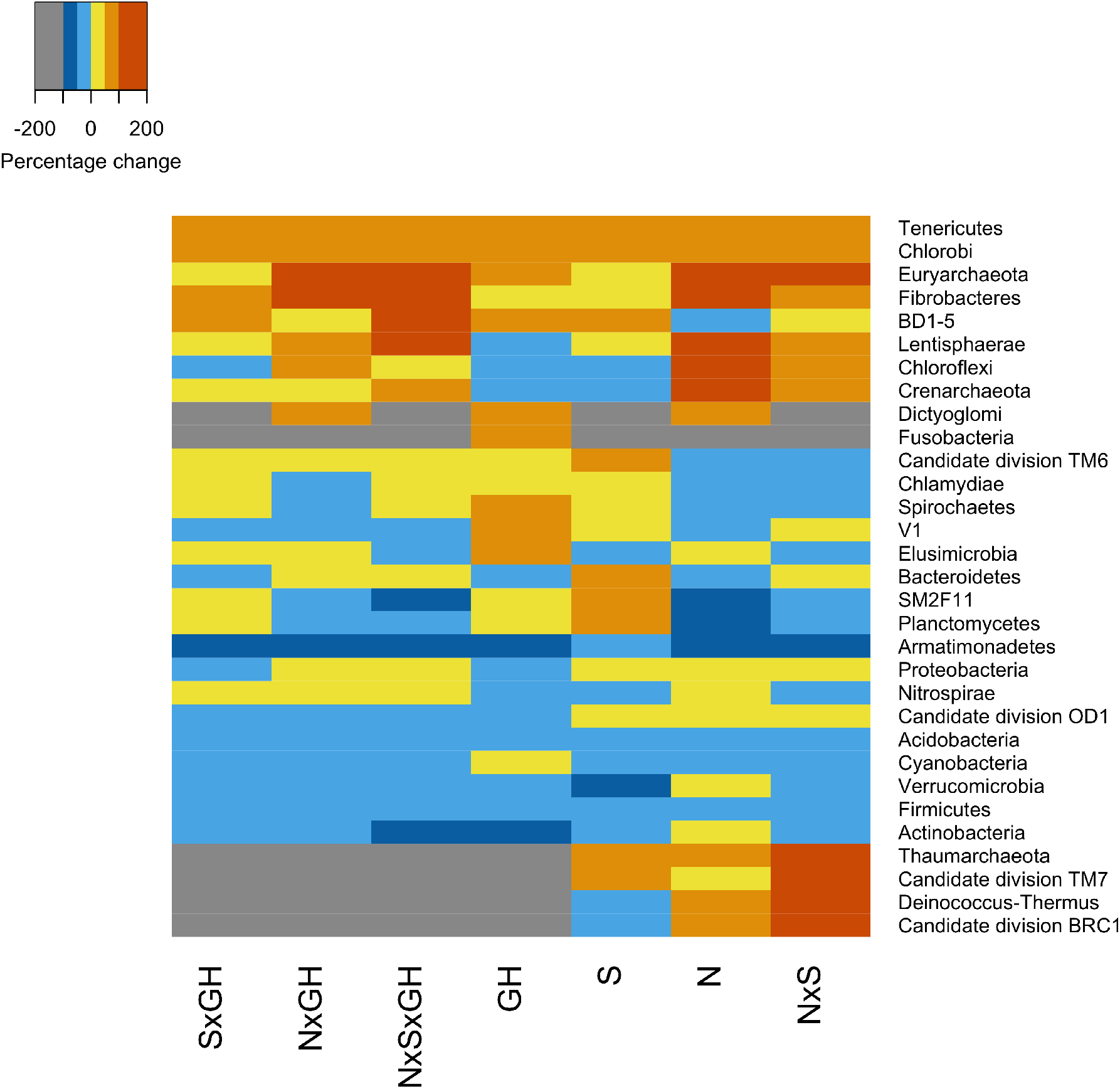
Relative change in the abundance of the 16S rRNA-derived phyla comparing the high level and the low level treatments, i.e. the relative abundance of each phylum at the high level was subtracted by the corresponding value for the low level and then divided with the low level value. Grey colour shows the phyla that disappeared with the high level treatment and red colour shows the phyla that appeared with the high level treatment. N, S and GH refer to the main treatment effects of nitrogen, sulfur and greenhouse, and NxS, NxGH and SxGH refer to their two-way interactions, while NxSxGH represents the three-way combination.

### Microbial metabolic traits

In addition to responses in the composition of operational taxonomic units and taxonomic groups at the phylum level, we assessed the treatment responses with regards to genome-encoded metabolic traits from shotgun metagenomic data. In total, protein-coding genes matching 4957 protein families (PFAMs) were found from the 31 metagenomes. Overall responses in the functional potential as determined by estimating Bray-Curtis distances revealed that only enhanced N deposition (high -N) caused a significant shift in functional attributes, and this was only observed below the water table (Table 1). There was also a significant interactive effect when warming and increased S deposition perturbations were combined (Table 1).

Moreover, we specifically searched for marker genes related to key steps in the anaerobic degradation of organic matter (e.g. nitrate- and sulfate reduction, hydrolysis, fermentation, methanogenesis) and aerobic methane and ammonia oxidation (Table S4). From 1057 possible responses at each depth horizon, merely 24 significant responses (p<0.05; FDR of 0.30) could be extracted from above the growing season mean water table (AWT), while the corresponding number in samples from below the growing season mean water table (BWT) was 30. At the AWT horizon, a few genes encoding for key steps in processes such as methanogenesis, sulfate reduction, nitrate reduction, sulfur oxidation, nitrogen fixation, syntrophy and hydrolysis, responded significantly to the experimental treatments (Fig. 3a). The majority of significant responses were connected to the 2-way interactive terms (NxS, NxGH and SxGH). Marker genes for methanogenesis significantly decreased in response to the main effects (N, S and GH), with amplification effects connected to the NxSxGH interaction, while they increased in response to the 2-way interactive terms. Genes encoding for proteins involved in dissimilatory sulfate reduction increased under enhanced S deposition, with amplification effects under the three-way perturbation and a decrease in the response when combined with warming or N.

**Fig. 3.**
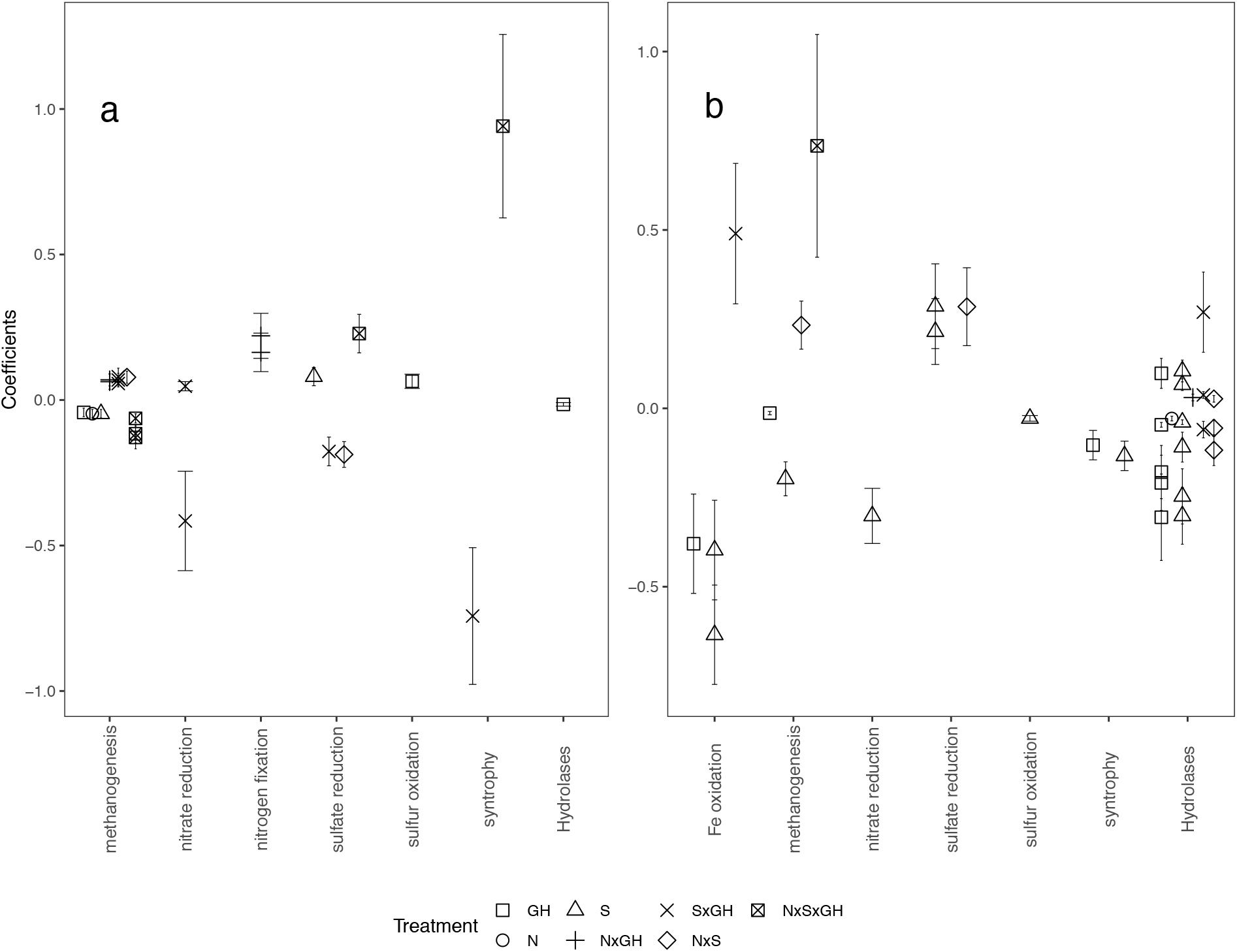
Generalized linear models (GLMs) on PFAMs (protein families) related to key steps in the anaerobic degradation of organic matter or relevant to the N and S cycling, at above the water table (AWT; panel a) and below the water table (BTW; panel b). The treatment responses are indicated by the coefficient, showing a decrease or increase in genome equivalents of each individual key PFAM. Only significant treatments responses are shown. Interaction terms in the regression models and the relationships among the variables in the model should be interpreted in as follows: positive coefficient in the case of interactions indicates a synergistic effect when combining perturbations, while a negative coefficient indicates an antagonistic effect, which can even lead to no expression of the perturbations.

At the BWT horizon, only a few genes encoding for methanogenesis, sulfate reduction, nitrate reduction, sulfur oxidation, syntrophy, and hydrolases responded significantly to the treatments. At this horizon the majority of the responses were connected to warming and enhanced S deposition, both resulting in a decrease of the different metabolic potentials, except for sulfate reduction that increased with the elevated S deposition (Fig. 3b). However, the few significant effects of the 2- and 3-way interactions with S or GH seemed to counteract these responses, resulting in an increase of the respective metabolic traits. For example, marker genes for methanogenesis, as observed for the AWT horizon, significantly decreased in response to the individual S and GH effects but increased in response to the NxS and NxSxGH perturbations. Genes encoding for hydrolases were the most affected among the studied metabolic traits, and mainly decreased with warming and enhanced S deposition as well as with simultaneous increase in N and S depositions.

### Co-variations between ecosystem functions and genetics in the light of perturbations

Taxonomic composition and genome-encoded traits were tightly coupled (p = 0.001, R = 0.87; Table S5). The previously reported realized methane production (Eriksson et al. 2010a), and the taxonomic composition data at 16S rRNA level were strongly related to each other across the different treatments and depth horizons (Table 2). However, genome-encoded functional traits as assessed from shotgun metagenomic data, including marker genes for methanogenesis and methanotrophy, did not correspond with measured function (i.e. methane production and oxidation; data no shown).

**Table 2.** Co-variation between taxonomic composition and process data (methane production and oxidation) assessed by fitting the process data onto an ordination derived from a nonmetric multidimensional scaling (NMDS), for every standardized depth (AWT: above the growing season mean water table, WT: around the growing season water table (WT), and BWT: below the growing season mean water table).

| Data | Standardized depth | NMDS stress value | R2 | Significance |
| --- | --- | --- | --- | --- |
| 16S rRNA - CH<sub>4</sub> production | AWT | 0.12 | 0.35 | 0.02 |
|  | WT | 0.09 | 0.39 | 0.017 |
|  | BWT | 0.06 | 0.51 | 0.004 |
| 16S rRNA gene -CH<sub>4</sub> production | AWT | 0.07 | 0.34 | 0.0026 |
|  | WT | 0.09 | 0.20 | ns |
|  | BWT | 0.12 | 0.30 | ns |
| 16S rRNA - CH<sub>4</sub> oxidation | AWT | 0.12 | 0.33 | ns |
|  | WT | 0.09 | 0.01 | ns |
|  | BWT | 0.06 | 0.43 | 0.016 |
| 16S rRNA gene -CH<sub>4</sub> oxidation | AWT | 0.07 | 0.05 | ns |
|  | WT | 0.09 | 0.31 | ns |
|  | BWT | 0.12 | 0.42 | 0.006 |

ns denotes non-significant results.

## Discussion

The long field experiment at the Degerö Stormyr peatland was established to investigate the effects of increased nitrogen (N) and sulfur (S) deposition as well as warming on methane and carbon dynamics in boreal oligotrophic mires (Granberg et al. 2001). Here we report on interactive long-term effects after 18 years of multiple perturbations on prokaryotic taxonomic diversity and genome-encoded traits, as well as their relationship with ecosystem-scale processes of interest *(i.e.* methane cycling, organic matter degradation and plant composition). The microbial taxonomic composition largely corresponds with results from previous studies of peatlands (Lin et al. 2012, Serkebaeva et al. 2013, Tveit et al. 2013). Thus, the proportion of archaeal sequences at 0.3% of total prokaryotic sequences was in the range of what has previously been observed for peat ecosystems (Tveit et al. 2013 and references therein). We also show that where vascular plants *(E. vaginatum, A. polifolia and V. oxycoccus)* replaced *Sphagnum* under N amendments (Wiedermann et al. 2007, Eriksson et al. 2010b), also prokaryotic communities shifted in composition in response to N additions. The concordance between beta-diversity of the plant vegetation composition derived from Wiedermann et al. (2007) and the composition of the peat prokaryote community, confirms the tight coupling between vegetation and microbes in the peat biome. The enrichments of roots at 10-15 cm into the peat vertical profile (Olid et al. 2017) likely play an essential role for this development by causing changes in organic matter composition and availability. The high level of N induced shifts in relative abundance among many phyla reported to harbour hydrolytic enzymes (Juottonen et al. 2017 and therein), i.e. increases in Proteobacteria and Actinobacteria, while Acidobacteria, Planctomycetes, and Bacteroidetes decreased. Interestingly, the decrease in Acidobacteria concomitantly with the relative increase in Proteobacteria may indicate a shift from the naturally nutrient poor conditions in the nutrient poor fen to more nutrient rich conditions because of the N amendment. In agreement with this, Lin et al. (2012) observed a higher abundance of Proteobacteria in rich fens compared to nutrient poor bogs. Since both phyla contain anaerobic carbohydrate polymer degrading representatives, these phyla may replace one another (cf. Schmidt et al. 2015) as result of changes in the type of carbohydrates supplied by the different plant communities forming the peat. The decrease in Cyanobacteria may reflect the decrease in *Sphagnum* abundance. Cyanobacteria occur in the hyalinic cell of the Sphagna lumena, where they have been shown to perform dinitrogen fixation (cf. Granhall & Selander 1973, Granhall & von Hofsten 1976, Berg et al. 2013). In addition to the taxonomic composition, also the genome-derived functional potential (PFAMs) was related to the composition of the vegetation. As such, our results fit the framework developed in studies of other soils subject to long-term N addition experiments, both with regards to microbial community composition and functional potential (Ramirez et al. 2010, Fierer et al. 2012, Leff et al. 2015, Zhou et al. 2015, Boot et al. 2016, Zeng et al. 2016), including grasslands, agricultural fields, agricultural black soils, and subalpine forests. For these systems, there is a consistent view that the shifts in microbial community composition and their metabolic functionality is due to the soil-plant-microbe interactions driven by N loading.

The significant relationships between prokaryotic community composition and methane oxidation and production, further emphasize the role of plant-prokaryotic interactions in regulating methane emissions. This is corroborated by the fact that all N amended to the plots is retained in the organic fraction of the peat (Eriksson 2010). However, responses in the overall set of genes, as well as the specific marker genes for methanogenesis and methanotrophy, did not match the patterns of observed methane production and consumption rates previously reported (Eriksson et al. 2010a). There are multiple possible explanations for the lack of correspondence, including an actual decoupling between gene abundance and their expression (Roling 2007, Freitag & Prosser 2009) or a high variability of the genomic content between individual samples. Also, temporal variation could play a role, as methane processing rates and genetic data were obtained during different years and locations within the treatment plots.

Similar to the elevated N deposition, also warming led to changes in prokaryotic community composition, supporting earlier findings of decreased methane production and emission rates while methane oxidation was unaffected (Eriksson et al. 2010a, Eriksson et al. 2010b). Thus, the relative abundance of genes involved in methanogenesis decreased in response to warming, while other metabolic traits such as methanotrophy were largely unaffected. The observed decrease in the methanogenic potential may be explained by lower input of easily degradable organic matter to the anoxic zone due to oxygen-exposure in the upper layers with higher temperature (Nilsson & Öquist 2009). Thus, the organic matter will be more recalcitrant when it is transferred into the permanent anoxic layer. In favour of this explanation, there was a decrease in the hydrolytic potential below the water table level pinpointing to a lower degradability of the biopolymers at these strata.

The enhanced S deposition affected the microbial taxonomic composition at the water table and the anoxic horizons. Although S amendment in the field did not have any effects on methane emissions, laboratory incubations of the methane producing-layers have shown a decrease in methane production by 55% in response to S amendments (Eriksson et al. 2010a, Eriksson et al. 2010b). This observation is supported by our results that the relative abundance of genes involved in methanogenesis were lower in the S-supplied plots. Below the water table, the observed decrease in hydrolytic and syntrophic potential combined with an increase of sulfate reduction potential, imply that organic matter degradation resulted in lower amounts of metabolic fermentation intermediates and H2 available for methanogenesis.

Above the water table, the sulfate reduction potential increased concomitantly with the observed decrease in methanogenic potential as expected from the thermodynamic constraints (Abram & Nedwell 1978, Kristjansson et al. 1982). Some of the sulfide generated in this process is likely emitted to the atmosphere at the prevailing low pH, while experimentally added S over the years has contributed to a 50% larger S-pool under ambient climate and to a ~15% larger S-pool when combined with the greenhouse treatment (Granberg et al. 2001, Åkerblom et al. 2013). A re-oxidation of this residual S-pool would result in a continuous supply of oxidized sulfur compounds that would sustain sulfate reduction in these treatments. The increase of the photosynthetic sulfur oxidizers, here represented by the phylum Chlorobi, in response to essentially all perturbations and in particular the S amendments, supports the presence of an internal sulfur cycle (Pester et al. 2010, Pester et al. 2012). Such a sulfur cycle is suggested to be involved in the regulation of the ratio between carbon dioxide to methane formation in peatlands (Pester et al. 2010, Pester et al. 2012).

The nitrogen applied has been shown to be completely retained for the duration of the experiment in the form of nitrogenous organic matter (Eriksson 2010). Because of this, the amendment of N to these highly nitrogen-limited systems is not expected to enhance the occurrence of ammonia oxidation, dissimilatory nitrate reduction to ammonia or denitrification. This is supported by the lack of any significant response of genes encoding for these processes in the N-amended plots. However, the 16S RNA analysis revealed that phyla hosting archaeal nitrifiers were present and increased in the plots with N addition. Especially, the positive response by groups within the Thaumarchaeota that have been shown to oxidize ammonia at very low levels, would potentially supply nitrite in the high-level N treatments. However, any nitrite formed would likely be immediately reduced by means of denitrification, anaerobic ammonium oxidation or assimilative or dissimilative reduction to ammonia.

## Conclusions

Experimental long-term treatments mimicking anthropogenic perturbations altered the microbial communities at the taxonomic level and to some extent redistribute genes encoding microbial metabolic profiles including changes in ecosystem-relevant traits, such as sulfate reduction and methanogenesis, partly coinciding with expressed overall ecosystem functions. The results from the 18-years field manipulation experiment emphasizes that interactive effects of multiple anthropogenic perturbations on ecosystem services lead to idiosyncratic and hard to predict disturbance-responses in natural microbial communities when studying each perturbation in isolation. The observed additive effects of the treatments on community composition and function emphasize the need for studying interactions among multiple anthropogenic perturbations to understand ecosystem responses to climate change.

## Acknowledgements

We thank Lucas Sinclair for bioinformatics support and the Uppsala Multidisciplinary Center for Advanced Computational Science (UPPMAX) for the computational and storage resources under projects b2014318. The project was mainly funded by the Swedish Research Council (contract no: 621-2011-4901) and additional research grants from the Swedish Research Council Formas to MN and SB and from the Swedish Research Council to AE, BS, MN and SB further supported the study. We are thankful to FEMS for the research fellowship (FRF 2014-1) award (MM).

