## Supplementary Material for "Effects of long-term warming and enhanced nitrogen and sulfur deposition on microbial communities in a boreal peatland"

[1] Department of Thematic Studies – Environmental Change, Linköping University, Linköping, Sweden
[2] Department of Clinical and Experimental Medicine, Linköping University, Linköping, Sweden
[3] Department of Ecology and Genetics, Limnology and Science for Life Laboratory, Uppsala University, Uppsala, Sweden
[4] eDNA solutions, Environmental DNA and bioinformatics solutions, Mölndal, Sweden
[5] Department of Biosciences, Center for BioGeoChemistry in the Anthropocene, Section for Aquatic Biology and Toxicology, University of Oslo, Oslo, Norway
[6] Department of Biology, Section of Microbiology, University of Copenhagen, Copenhagen, Denmark
[7] Department of Clinical Laboratory Sciences, Jouf University, Qurayyat, Saudi Arabia
[8] Department of Forest Ecology & Management, Swedish University of Agricultural Sciences, Umeå, Sweden

**Table S1** Distribution of samples withdrawn from the long-term experimental field at Degerö Stormyr for the microbial characterization carried out in the present study. S: sulfate treatment, N: nitrogen treatment, GH: warming treatment. A - E: sampled depths. Light grey, dark grey, and black: depths classified as above-, around-, and below the growing season mean water table levels, respectively. 16S shows the samples analyzed for 16S rRNA and 16S rRNA gene. PFAM: Protein FAMilies denote samples targeted by shotgun metagenome sequencing (31 samples). * marks samples not included in the analyses (one 16S rRNA and one PFAM sample). [d] marks 16S rRNA gene samples excluded from the alpha-diversity analysis due to low total number of sequences. [r] marks 16S rRNA samples excluded from the alpha-diversity analysis due to low total number of sequences.

| Treatment | Plot | A (9cm) | B (13cm) | C (17cm) | D (21cm) | E (25cm) |
|---|---|---|---|---|---|---|
| Control | 11 | 16S | 16S;PFAM | 16S | 16S;PFAM | 16S |
| Control | 19 | 16S | 16S;PFAM | 16S | 16S;PFAM | 16S |
| GH | 4 | 16S | 16S;PFAM | 16S | 16S;PFAM | 16S |
| GH | 16 | 16S | 16S;PFAM | 16S | 16S;PFAM | 16S |
| N | 12 | 16S | 16S;PFAM | 16S | 16S;PFAM | 16S |
| N | 18 | 16S[r] | 16S;PFAM | 16S | 16S;PFAM | 16S* |
| NxGH | 9 | 16S | 16S;PFAM | 16S | 16S;PFAM | 16S |
| NxGH | 17 | 16S | 16S;PFAM | 16S | 16S;PFAM | 16S |
| NxS | 8 | 16S | 16S;PFAM | 16S | 16S;PFAM | 16S |
| NxS | 14 | 16S | 16S;PFAM | 16S | 16S;PFAM | 16S |
| NxSxGH | 2 | 16S[d] | 16S;PFAM | 16S | 16S;PFAM | 16S |
| NxSxGH | 13 | 16S | 16S;PFAM | 16S | 16S;PFAM | 16S |
| S | 3 | 16S | 16S[d];PFAM | 16S | 16S;PFAM | 16S |
| S | 7 | 16S | 16S;PFAM | 16S | 16S;PFAM | 16S |
| SxGH | 5 | 16S | 16S;PFAM | 16S | 16S;PFAM | 16S[r] |
| SxGH | 10 | 16S | 16S;PFAM* | 16S | 16S;PFAM | 16S |

**Table S2** Evaluation matrix for treatments effects. Low and high refer to the level of the treatments: for sulfate low level: ambient (3 kg) or high level amended by $Na_2SO_4$ to reach 20 kg S ha$^{-1}$ yr$^{-1}$, for nitrogen low level: ambient (2 kg) or high level addition of $NH_4NO_3$ to reach 30 kg N ha$^{-1}$ yr$^{-1}$ and for warming high level: greenhouse (GH) cover or low level: ambient conditions. AWT: above the growing season mean water table level. WT: around the growing season mean water table level. BWT: below the growing season mean water table level. * sample that could not be included in the PFAM analysis at AWT, thus one replicate less was included from what is displayed in the Factorial design.

| Plot number | Field treatment | Number of replicates | | | Main effect | | | | | | 2-way interaction effect | | | | | | 3-way interaction effect | |
|---|---|---|---|---|---|---|---|---|---|---|---|---|---|---|---|---|---|---|
| | | | | | N | | S | | GH | | NxS | | NxGH | | SxGH | | NxSxGH | |
| | | AWT | WT | BWT | low | high | low | high | low | high | low | high | low | high | low | high | low | high |
| 11 | control | 1 | 1 | 3 | x | | x | | x | | x | | x | | x | | x | |
| 19 | control | 1 | 1 | 3 | x | | x | | x | | x | | x | | x | | x | |
| 4 | GH | 1 | 1 | 3 | x | | x | | | x | x | | | | | | | |
| 16 | GH | 1 | 1 | 3 | x | | x | | | x | x | | | | | | | |
| 12 | N | 1 | 2 | 2 | | x | x | | x | | | | | | x | | | |
| 18 | N | 1 | 2 | 1 | | x | x | | x | | | | | | x | | | |
| 9 | NxGH | 1 | 1 | 3 | | x | x | | | x | | | | x | | | | |
| 17 | NxGH | 1 | 1 | 3 | | x | x | | | x | | | | x | | | | |
| 8 | NxS | 2 | 2 | 1 | | x | | x | x | | | x | | | | | | |
| 14 | NxS | 1 | 1 | 3 | | x | | x | x | | | x | | | | | | |
| 2 | NxSxGH | 2 | 1 | 2 | | x | | x | | x | | x | | x | | x | | x |
| 13 | NxSxGH | 1 | 1 | 3 | | x | | x | | x | | x | | x | | x | | x |
| 3 | S | 1 | 1 | 3 | x | | | x | x | | | | x | | | | | |
| 7 | S | 1 | 1 | 3 | x | | | x | x | | | | x | | | | | |
| 5 | SxGH | 1 | 1 | 3 | x | | | x | | x | | | | | | x | | |
| 10 | SxGH* | 1 | 1 | 3 | x | | | x | | x | | | | | | x | | |
| Statistical replicates | Factorial design | | | | 8 | 8 | 8 | 8 | 8 | 8 | 4 | 4 | 4 | 4 | 4 | 4 | 2 | 2 |
| | AWT | | | | 8 | 10 | 8 | 10 | 9 | 9 | 4 | 6 | 4 | 5 | 4 | 5 | 2 | 3 |
| | WT | | | | 8 | 11 | 10 | 9 | 11 | 8 | 4 | 5 | 4 | 4 | 6 | 4 | 2 | 2 |
| | BWT | | | | 24 | 18 | 21 | 21 | 19 | 23 | 12 | 9 | 12 | 11 | 9 | 11 | 6 | 5 |

**Table S3** Depth normalization based on the plot specific average value as measured in the years 2004-2006 (Eriksson, et al., 2010a). MTW05: mean water table level in 2005 at each plot. An average of the MTW05 over all plots was calculated (8.8 cm). Difference: the MTW05 of each plot subtracted from the average (8.8 cm). Adjusted water level: difference added to 14 cm (corresponding to the 2004 – 2006 mean growing season water table level). A – E: sampled depths classified to above the growing season mean water table level (AWT), around the growing season mean water table level (WT), and below the growing season mean water level (BWT).

| Treatment | Plot | MTW05 (cm) | Difference (cm) | Adjusted water level | A (7-11cm) | B (11-15cm) | C (15-19cm) | D (19-23cm) | E (23-27cm) |
|---|---|---|---|---|---|---|---|---|---|
| Control | 11 | 9.1 | -0.4 | 13.7 | AWT | WT | BWT | BWT | BWT |
| Control | 19 | 14.8 | -6.0 | 8.0 | AWT | WT | BWT | BWT | BWT |
| GH | 4 | 10.0 | -1.2 | 12.8 | AWT | WT | BWT | BWT | BWT |
| GH | 16 | 10.8 | -2.0 | 12.0 | AWT | WT | BWT | BWT | BWT |
| N | 12 | 7.7 | 1.1 | 15.1 | AWT | WT | WT | BWT | BWT |
| N | 18 | 7.9 | 0.8 | 14.8 | AWT | WT | WT | BWT | BWT |
| NxGH | 9 | 7.1 | 1.7 | 15.7 | AWT | WT | WT | BWT | BWT |
| NxGH | 17 | 10.0 | -1.2 | 12.8 | AWT | WT | BWT | BWT | BWT |
| NxS | 8 | 3.6 | 5.2 | 19.2 | AWT | AWT | WT | WT | BWT |
| NxS | 14 | 9.7 | -0.9 | 13.1 | AWT | WT | BWT | BWT | BWT |
| NxSxGH | 2 | 5.7 | 3.1 | 17.1 | AWT | AWT | WT | BWT | BWT |
| NxSxGH | 13 | 10.4 | -1.7 | 12.3 | AWT | WT | BWT | BWT | BWT |
| S | 3 | 9.3 | -0.5 | 13.5 | AWT | WT | BWT | BWT | BWT |
| S | 7 | 8.7 | 0.1 | 14.1 | AWT | WT | BWT | BWT | BWT |
| SxGH | 5 | 12.0 | -3.3 | 10.7 | AWT | WT | BWT | BWT | BWT |
| SxGH | 10 | 10.1 | -1.3 | 12.7 | AWT | WT | BWT | BWT | BWT |

**Table S4** Selected marker genes related to the key steps in the anaerobic degradation of organic matter as well as relevant to the nitrogen and sulfur cycling. The Protein FAMilies (PFAM) that were significant to at least one treatment, are marked in bold.

| PFAM | Description | Metabolic pathway |
|---|---|---|
| PF02461 | ammonia monooxygenase | ammonia oxidation I (aerobic) |
| PF04744.1 | ammonia monooxygenase | ammonia oxidation I (aerobic) |
| PF04896.1 | ammonia monooxygenase | ammonia oxidation I (aerobic) |
| PF13435 | cytochrome c554 | ammonia oxidation IV (autotrophic ammonia oxidizers) |
| PF14100.1 | particulate methane monooxygenase hydroxylase component | ammonia oxidation I (aerobic) |
| PF00032 | cytochrome bc1 | Fe(II) oxidation |
| PF02335 | cytochrome c-552 Fe and ammonium oxidation | Fe(II) oxidation |
| PF13631 | cytochrome bc1 | Fe(II) oxidation |
| PF00056.18 | lactate/malate dehydrogenase, NAD binding domain | L-lactate dehydrogenase |
| PF02615.9 | Malate/L-lactate dehydrogenase | L-lactate dehydrogenase |
| PF02866.13 | lactate/malate dehydrogenase, alpha/beta C-terminal domain | L-lactate dehydrogenase |
| PF00763 | Tetrahydrofolate dehydrogenase/cyclohydrolase, catalytic domain | homoacetogenesis |
| PF02882 | PF13631 | homoacetogenesis |
| **PF00871** | **acetate kinase** | **methanogenesis_acetate** |
| PF01493 | formylmethanofuran dehydrogenase, tungsten enzyme | methanogenesis_co2 |
| PF01568 | formylmethanofuran dehydrogenase | methanogenesis_co2 |
| PF01913 | formylmethanofuran:H4SPT formyltransferase | methanogenesis_co2 |
| PF01993 | F420-dependent methylene-H4MPT reductase | methanogenesis_co2 |
| PF02007 | methyl-H4MPT:coenzyme M methyltransferase | methanogenesis_co2 |
| **PF02240** | **Methyl-coenzyme M reductase operon protein G** | **methanogenesis** |
| PF02241 | Methyl-coenzyme M reductase operon protein B | methanogenesis |
| PF02505 | Methyl-coenzyme M reductase operon protein D | methanogenesis |
| **PF02663** | **formylmethanofuran dehydrogenase** | **methanogenesis_co2** |
| PF02741 | formylmethanofuran:H4SPT formyltransferase | methanogenesis_co2 |
| PF02783 | Methyl-coenzyme M reductase operon protein B | methanogenesis |
| PF03598 | acetyl-CoA decarbonylase/synthase complex _ subunit | methanogenesis_acetate |
| PF03599 | acetyl-CoA decarbonylase/synthase complex __ component | methanogenesis_acetate |

| | | |
|---|---|---|
| PF04207 | methyl-H4MPT:coenzyme M methyltransferase | methanogenesis_co2 |
| PF04208 | methyl-H4MPT:coenzyme M methyltransferase | methanogenesis_co2 |
| PF04210 | methyl-H4MPT:coenzyme M methyltransferase | methanogenesis_co2 |
| PF04211 | methyl-H4MPT:coenzyme M methyltransferase | methanogenesis_co2 |
| **PF04609** | **Methyl-coenzyme M reductase operon protein C** | **methanogenesis** |
| PF05440 | methyl-H4MPT:coenzyme M methyltransferase | methanogenesis_co2 |
| **PF09472** | **methyl-H4MPT:coenzyme M methyltransferase** | **methanogenesis_co2** |
| PF02406 | soluble methane monooxygenase | methane oxidation to methanol I |
| PF04744.2 | particulate methane monooxygenase hydroxylase component | methane oxidation to methanol II |
| PF04896.2 | particulate methane monooxygenase hydroxylase component | methane oxidation to methanol II |
| PF14100.2 | particulate methane monooxygenase hydroxylase component | methane oxidation to methanol II |
| **PF00420** | **NADH:quinone oxidoreductase I** | **nitrate reduction VIII (dissimilatory)** |
| PF00499 | NADH:quinone oxidoreductase I | nitrate reduction VIII (dissimilatory) |
| PF00795 | assimilatory nitrite reductase | nitrate reduction V (assimilatory) |
| PF02613 | respiratory nitrate reductase | nitrate reduction I (denitrification) |
| PF02665 | respiratory nitrate reductase | nitrate reduction I (denitrification) |
| **PF03892** | **periplasmic nitrate reductase** | **nitrate reduction X (periplasmic, dissimilatory)** |
| **PF03927** | **periplasmic nitrate reductase** | **nitrate reduction X (periplasmic, dissimilatory)** |
| PF05088 | glutamate dehydrogenase (NADP+) | nitrate reduction V (assimilatory) |
| PF09163.2 | formate dehydrogenase N | nitrate reduction III (dissimilatory) |
| PF00142.13 | 4Fe-4S iron sulfur cluster binding proteins, NifH/frxC family | nitrogen fixation |
| PF00142 | [FeFe]-nitrogenase complex | nitrogen fixation (flavodoxin) |
| PF02579.12 | Dinitrogenase iron-molybdenum cofactor | nitrogen fixation |
| PF02579 | Dinitrogenase iron-molybdenum cofactor | nitrogen fixation I (ferredoxin) |
| PF03139 | [FeFe]-nitrogenase complex | nitrogen fixation II (flavodoxin) |
| **PF03206.9** | **Nitrogen fixation protein NifW** | **nitrogen fixation** |
| PF04319.8 | NifZ domain | nitrogen fixation |
| **PF04891.7** | **NifQ** | **nitrogen fixation** |
| PF01077 | assimilatory sulfite reductase/nitrite reductase (dissimilatory) | sulfate reduction I (assimilatory) |
| PF01078 | dissimilatory sulfite reductase | sulfate reduction V (dissimilatory) |
| PF01507 | phosphoadenosine phosphosulfate reductase | sulfate reduction I (assimilatory) |
| **PF01583** | **adenylylsulfate kinase** | **sulfate reduction I (assimilatory)** |

| | | |
|---|---|---|
| PF03460 | assimilatory sulfite reductase/nitrite reductase (dissimilatory) | sulfate reduction I (assimilatory) |
| **PF03461** | **dissimilatory sulfite reductase** | **sulfate reduction V (dissimilatory)** |
| PF03464 | DsrK DsrC-disulfide reductase | sulfate reduction IV (dissimilatory) |
| PF03465 | dissimilatory sulfite reductase | sulfate reduction IV (dissimilatory) |
| PF03466 | dissimilatory sulfite reductase | sulfate reduction V (dissimilatory) |
| PF00581 | rhodanese 2599 | superpathway of sulfide oxidation (phototrophic sulfur bacteria) |
| PF01206 | TusA sulfur-carrier protein | sulfur oxidation IV (intracellular sulfur) |
| PF02635 | DsrEFH hexamer | sulfur oxidation IV (intracellular sulfur) |
| **PF04358** | **DsrC dimer** | **sulfur oxidation IV (intracellular sulfur)** |
| PF13686 | DsrEFH hexamer | sulfur oxidation IV (intracellular sulfur) |
| PF00766 | Electron transfer flavoprotein FAD-binding domain | Electron transfer flavoprotein-ubiquinone oxidoreductase |
| **PF01012** | **Electron transfer flavoprotein domain** | **Electron transfer flavoprotein-ubiquinone oxidoreductase** |
| PF02634 | FdhD/NarQ family | membranebound FDH |
| **PF04216** | **Protein involved in formate dehydrogenase formation** | **FDH** |
| PF04422 | Coenzyme F420 hydrogenase/dehydrogenase, beta subunit N-term | FDH |
| PF04432 | Coenzyme F420 hydrogenase/dehydrogenase, beta subunit C terminus | FDH |
| PF05187 | Electron transfer flavoprotein-ubiquinone oxidoreductase | Electron transfer flavoprotein-ubiquinone oxidoreductase |
| PF09163.1 | Formate dehydrogenase N, transmembrane | membranebound FDH |
| PF10125 | NADH dehydrogenase I, subunit N related protein | membranebound H2ase |
| PF10588 | NADH-ubiquinone oxidoreductase-G iron-sulfur binding region | FeS oxidoreductase |
| PF10589 | NADH-ubiquinone oxidoreductase-F iron-sulfur binding region | FeS oxidoreductase |
| PF11390 | NADH-dependant formate dehydrogenase delta subunit FdsD | NADH-linked FDH |
| PF14498.1 | Glycosyl hydrolase family 65, N-terminal domain | Hydrolysis |
| PF14587.1 | O-Glycosyl hydrolase family 30 | Hydrolysis |
| PF00232.13 | Glycosyl hydrolase family 1 | Hydrolysis |
| PF00331.15 | Glycosyl hydrolase family 10 | Hydrolysis |
| PF12899.2 | Alkaline and neutral invertase | Hydrolysis |
| PF05838.7 | Glycosyl hydrolase 108 | Hydrolysis |
| PF00457.12 | Glycosyl hydrolases family 11 | Hydrolysis |
| PF03537.8 | Glycoside-hydrolase family GH114 | Hydrolysis |

| | | |
|---|---|---|
| PF01670.11 | Glycosyl hydrolase family 12 | Hydrolysis |
| PF01373.12 | Glycosyl hydrolase family 14 | Hydrolysis |
| PF00723.16 | Glycosyl hydrolases family 15 | Hydrolysis |
| PF00722.16 | Glycosyl hydrolases family 16 | Hydrolysis |
| PF00332.13 | Glycosyl hydrolases family 17 | Hydrolysis |
| PF00704.23 | Glycosyl hydrolases family 18 | Hydrolysis |
| PF00182.14 | Chitinase class I | Hydrolysis |
| PF00703.16 | Glycosyl hydrolases family 2 | Hydrolysis |
| PF02836.12 | Glycosyl hydrolases family 2, TIM barrel domain | Hydrolysis |
| PF02837.13 | Glycosyl hydrolases family 2, sugar binding domain | Hydrolysis |
| PF00728.17 | Glycosyl hydrolase family 20, catalytic domain | Hydrolysis |
| PF02838.10 | Glycosyl hydrolase family 20, domain 2 | Hydrolysis |
| PF01183.15 | Glycosyl hydrolases family 25 | Hydrolysis |
| PF02156.10 | Glycosyl hydrolase family 26 | Hydrolysis |
| PF00295.12 | Glycosyl hydrolases family 28 | Hydrolysis |
| PF00933.16 | Glycosyl hydrolase family 3 N terminal domain | Hydrolysis |
| PF01915.17 | Glycosyl hydrolase family 3 C-terminal domain | Hydrolysis |
| PF02055.11 | O-Glycosyl hydrolase family 30 | Hydrolysis |
| PF01055.21 | Glycosyl hydrolases family 31 | Hydrolysis |
| PF08244.7 | Glycosyl hydrolases family 32 C terminal | Hydrolysis |
| PF00251.15 | Glycosyl hydrolases family 32 N-terminal domain | Hydrolysis |
| PF01301.14 | Glycosyl hydrolases family 35 | Hydrolysis |
| PF01074.17 | Glycosyl hydrolases family 38 N-terminal domain | Hydrolysis |
| PF07748.8 | Glycosyl hydrolases family 38 C-terminal domain | Hydrolysis |
| PF01229.12 | Glycosyl hydrolases family 39 | Hydrolysis |
| PF02056.11 | Family 4 glycosyl hydrolase | Hydrolysis |
| PF02449.10 | Beta-galactosidase | Hydrolysis |

| | | |
|---|---|---|
| PF08533.5 | Beta-galactosidase C-terminal domain | Hydrolysis |
| PF08532.5 | Beta-galactosidase trimerisation domain | Hydrolysis |
| PF04616.9 | Glycosyl hydrolases family 43 | Hydrolysis |
| PF12891.2 | Glycoside hydrolase family 44 | Hydrolysis |
| PF01532.15 | Glycosyl hydrolase family 47 | Hydrolysis |
| PF02011.10 | Glycosyl hydrolase family 48 | Hydrolysis |
| PF03718.8 | Glycosyl hydrolase family 49 | Hydrolysis |
| PF11975.3 | Family 4 glycosyl hydrolase C-terminal domain | Hydrolysis |
| PF03512.8 | Glycosyl hydrolase family 52 | Hydrolysis |
| PF07745.8 | Glycosyl hydrolase family 53 | Hydrolysis |
| PF03065.10 | Glycosyl hydrolase family 57 | Hydrolysis |
| PF02057.10 | Glycosyl hydrolase family 59 | Hydrolysis |
| PF01341.12 | Glycosyl hydrolases family 6 | Hydrolysis |
| PF03664.8 | Glycosyl hydrolase family 62 | Hydrolysis |
| PF03200.11 | Mannosyl oligosaccharide glucosidase | Hydrolysis |
| PF03633.10 | Glycosyl hydrolase family 65, C-terminal domain | Hydrolysis |
| PF03632.10 | Glycosyl hydrolase family 65 central catalytic domain | Hydrolysis |
| PF03636.10 | Glycosyl hydrolase family 65, N-terminal domain | Hydrolysis |
| PF13199.1 | Glycosyl hydrolase family 66 | Hydrolysis |
| PF07477.7 | Glycosyl hydrolase family 67 C-terminus | Hydrolysis |
| PF07488.7 | Glycosyl hydrolase family 67 middle domain | Hydrolysis |
| PF03648.9 | Glycosyl hydrolase family 67 N-terminus | Hydrolysis |
| PF03659.9 | Glycosyl hydrolase family 71 | Hydrolysis |
| PF03198.9 | Glucanosyltransferase | Hydrolysis |
| PF03663.9 | Glycosyl hydrolase family 76 | Hydrolysis |
| PF02446.12 | 4-alpha-glucanotransferase | Hydrolysis |
| PF03662.9 | Glycosyl hydrolase family 79, N-terminal domain | Hydrolysis |
| PF01270.12 | Glycosyl hydrolases family 8 | Hydrolysis |
| PF03639.8 | Glycosyl hydrolase family 81 | Hydrolysis |
| PF03644.8 | Glycosyl hydrolase family 85 | Hydrolysis |
| PF07470.8 | Glycosyl Hydrolase Family 88 | Hydrolysis |
| PF00759.14 | Glycosyl hydrolase family 9 | Hydrolysis |
| PF07971.7 | Glycosyl hydrolase family 92 | Hydrolysis |

| | | |
|---|---|---|
| PF10566.4 | Glycoside hydrolase 97 | Hydrolysis |
| PF11790.3 | Glycosyl hydrolase catalytic core | Hydrolysis |

**Table S5** Correspondence analysis between prokaryotic community composition, vegetation community composition (Wiedermann, et al., 2007) and functional potential, as assessed by procrustes superimposition analyses, for the standardized depth (AWT: above the growing season mean water table, WT: around the growing season mean water table (WT), and BWT: below the growing season mean water table).

| Data | Standardized depth | Procrustes Sum of Squares (m12 squared) | Correlation in a symmetric Procrustes rotation | Significance |
|---|---|---|---|---|
| 16S rRNA - vegetation | AWT | 0.596 | 0.6356 | 0.002 |
| | WT | 0.5812 | 0.6471 | 0.001 |
| | BWT | 0.7389 | 0.511 | 0.001 |
| 16S rRNA gene - vegetation | AWT | 0.5398 | 0.6784 | 0.001 |
| | WT | 0.5969 | 0.6349 | 0.001 |
| | BWT | 0.8133 | 0.432 | 0.001 |
| Pfam - vegetation | AWT | 0.7694 | 0.4803 | 0.281 |
| | BWT | 0.58 | 0.648 | 0.007 |
| 16S rRNA - 16S rRNA gene | AWT | 0.0504 | 0.9745 | 0.001 |
| | WT | 0.0389 | 0.9804 | 0.001 |
| | BWT | 0.08112 | 0.9586 | 0.001 |
| 16S rRNA - Pfam | AWT | 0.5223 | 0.6912 | 0.501 |
| | BWT | 0.5012 | 0.7062 | 0.149 |
| 16S rRNA gene - Pfam | AWT | 0.24 | 0.8718 | 0.001 |
| | BWT | 0.2493 | 0.8664 | 0.001 |

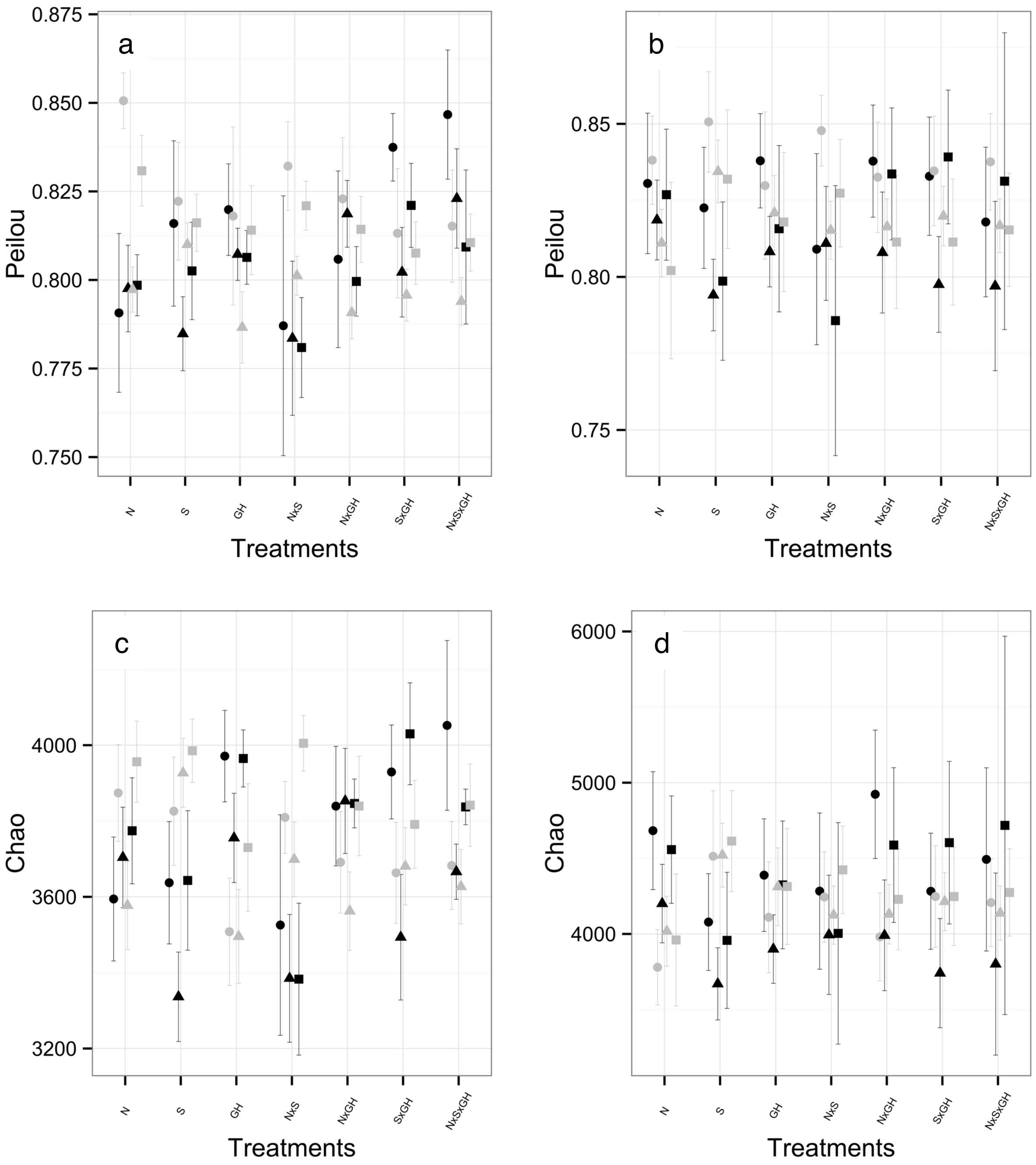


**Fig. S1** Alpha diversity of the prokaryotic community derived from the 16 rRNA gene (a, c) and derived from the 16 rRNA (b, d), assessed by richness estimate Pielous index (a, b) and evenness estimate Chao (c, d). The high levels of the treatments are shown in black and the low levels in grey. The circle, square and triangle represent the above the growing season mean water table level, around the growing season mean water level and below the growing season mean water table level depth horizons, respectively.

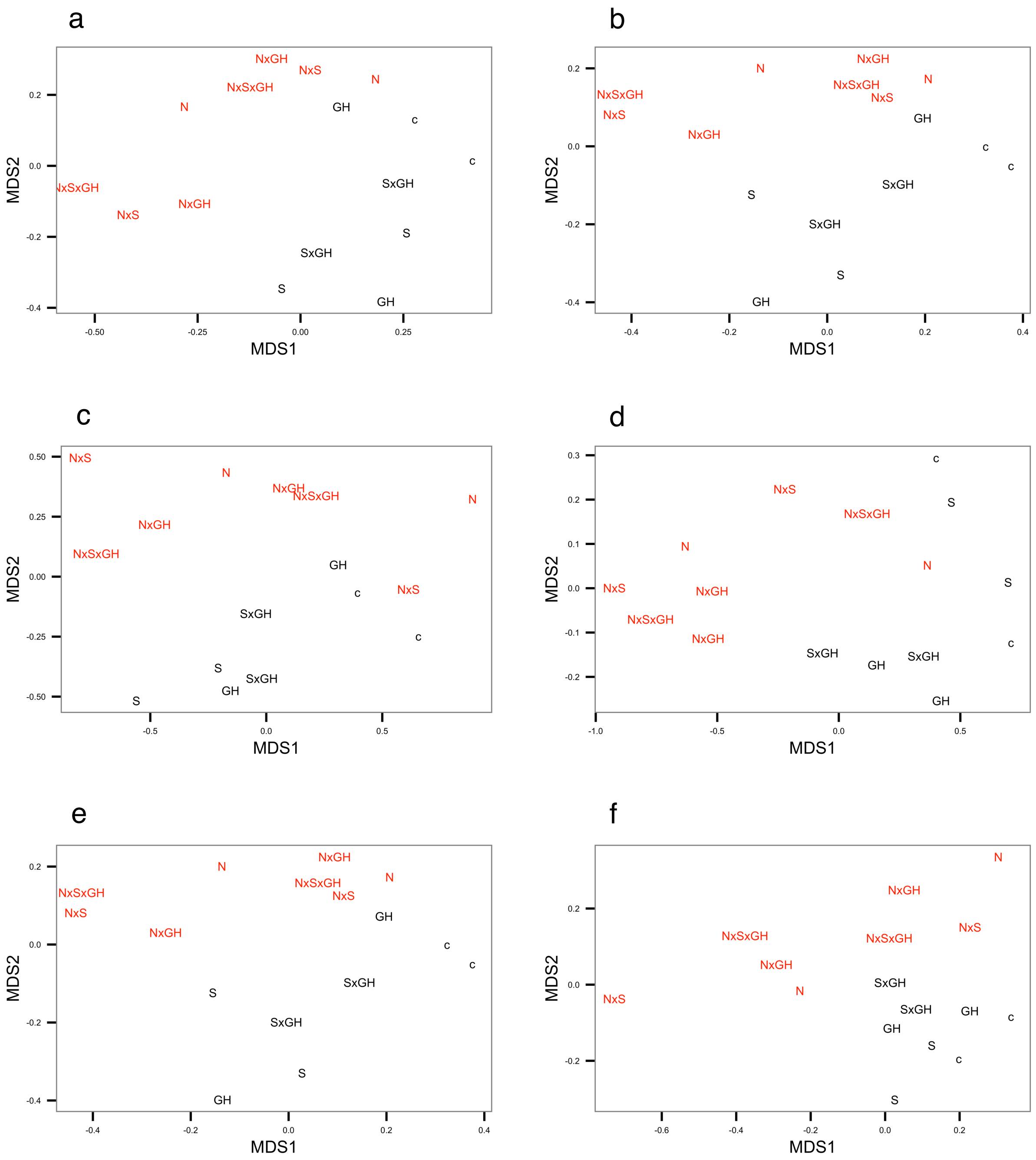


**Fig. S2** Non-metric multidimensional scaling plot of the microbial community composition, derived from 16S rRNA (a-c) and 16S rRNA gene (D-E), among the treatment plots and separated by the depths horizons at above the growing season mean water table level (a, d), around the growing season mean water table level (b, e) and below the growing season mean water table level (c, f). For the stress values see the article Table 2. The treatments receiving nitrogen are represented in red and the treatments without receiving nitrogen are represented in black.

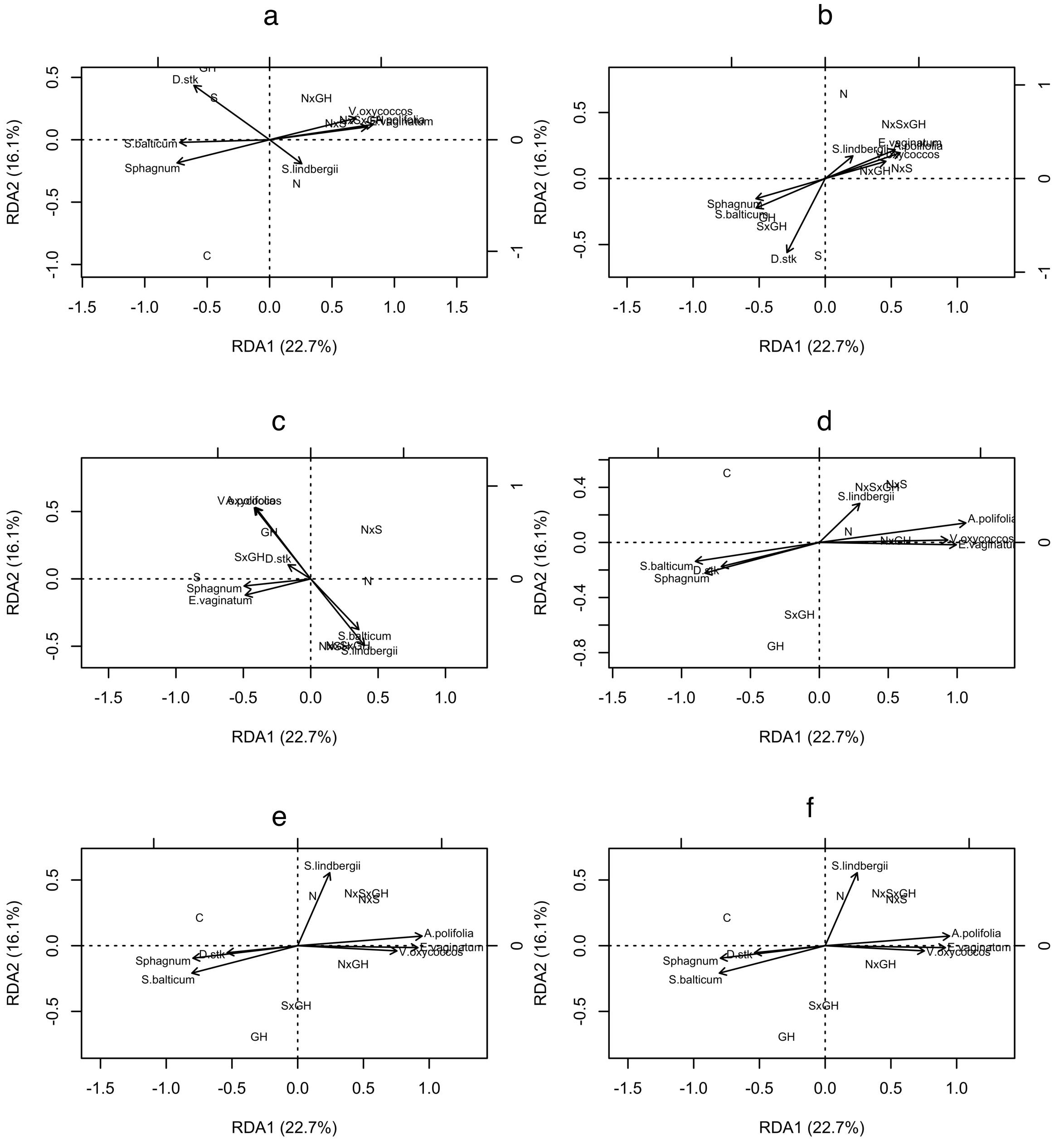


**Fig. S3** Distance-based Redundancy analysis of the prokaryote community in relation to the vegetation community, derived from Wiedermann *et al*., (2007), at above the growing season mean water table level (a, d), around the growing season mean water table level (b, e) and below the growing season mean water table level (c, f). A to C: OTUs derived from the 16S rRNA gene. D to F: OTUs derived from the 16S rRNA.
